# Predicting the Effects of Hemlock Woolly Adelgid on Microhabitat Structure and Small Mammal Communities

**DOI:** 10.1101/269142

**Authors:** Allyson L. Degrassi

## Abstract

Hemlock woolly adelgid (HWA) invasion and preemptive logging practices alter the habitat structure of New England forests and may indirectly affect associated small mammal communities. Microhabitat structure was measured and small mammals were censused in eight large experimental plots to quantify and predict these effects. The Harvard Forest Long-Term Ecological Research experiment is a replicated two-block design that includes four 0.81-ha canopy treatments: 1) Hemlock Control, 2) Hardwood Control, 3) Girdled Treatment, in which hemlock trees were killed by girdling in 2005 and left standing to simulate HWA invasion, and 4) Logged Treatment, in which trees were removed to simulate preemptive logging management practices. Nine microhabitat characteristics were measured from plot photos revealing differences among microhabitat structure. Small mammals were censused with arrays of 49 Sherman traps per plot and population sizes of common species were estimated with mark-recapture analysis. Between 6 and 8 mammal species were recorded in all treatments and species composition varied slightly. Populations of two common rodents (*Peromyscus spp*.) were not affected by treatment, but the southern red-backed vole population was greatest in the Girdled treatment. Estimated species richness was greater in the Girdled treatment than the Hemlock control, but richness did not differ between Girdled and Logged treatments, which suggests preemptive logging is as detrimental to some small mammal species as HWA invasion. Overall, nine years post disturbance, there is little evidence of a major shift in small mammal community structure in response to woolly adelgid invasion, with only minor changes in relative abundance.

## Introduction

Eastern hemlock (*Tsuga canadensis* (L.) Carrière) is considered a foundation species because it is an abundant primary producer with direct and indirect links to many other species in its food web (Ellison et al. 2005a). Their populations have declined dramatically in the eastern United States because of the damage inflicted on them by the non-native invasive insect (McClure 1991, Orwig and Foster 1998, Orwig et al. 2002, Ellison et al. 2005a), hemlock woolly adelgid (*Adelges tsugae*, Annand 1928) which is native to Asia and was introduced to the United States in the 1950’s (McClure 1989). It is a sap sucking insect that defoliates trees (Orwig et al. 2008) and causes rapid hemlock mortality (McCulre 1991). The damage to individual trees spreads through forests and decreases canopy cover, which alters the forest by increasing the amount of light that reaches the forest floor and consequently promoting new growth of hardwood species (Farnsworth et al. 2012). Because the hemlock woolly adelgid threatens much of the old growth forests in the eastern United States, forest management strategies such as preemptive logging were being considered to slow the spread of the adelgid and to conserve late successional forests (Foster and Orwig 2006).

Dramatic changes in forest structure caused by HWA invasion can shift ecosystem processes and shifts in biodiversity. These shifts within the ecosystem affect taxonomic groups differently. For example, loss of eastern hemlocks results in an increase in local ant species diversity (Ellison et al. 2005b), but a decrease in regional bird (Tingley et al. 2002) and salamander (Siddig et al. 2016) population and occurrence. These inconsistent responses of animal diversity to hemlock loss over varying taxonomic groups make it difficult to predict how species will prevail after the loss of hemlocks, which are not expected to recover from hemlock woolly adelgid invasion. This imminent loss of eastern hemlock foundation species will cause ecosystem changes within the forest, but how this change in habitat structure will impact supported species distribution is generally unknown and may be different depending on the species in question (Table 1).

**Table 1.** Hypothesized influence of simulated HWA on relative abundance of individuals within small mammal communities at Harvard Forest in Petersham, MA.

| Small Mammal Species | Predicted direction of influence of<br>HWA on small mammal abundance<br>relative to Hemlock control<br>(↑ increase, ↓ decrease, 0 no effect) | Supporting Literature |
| --- | --- | --- |
| Deer mouse<br>*Peromyscus maniculatus* | 0 | Graves et al. 1988 |
| White-footed mouse<br>*Peromyscus leucopus* | ↓ | Henein et al. 1998; Graves<br>et al. 1988 |
| Woodland jumping mouse<br>*Napaeozapus insignis* | ↑ | Vickery and Rivest 1992 |
| Southern red-backed vole<br>*Myodes gapperi* | ↓ | Merritt 1981 |
| Southern flying squirrel<br>*Glaucomys volans* | ↑ | Taulman and Seaman 2000 |
| American red squirrel<br>*Tamiasciurus hudsonicus* | ↑ | Ransome et al. 1997 |
| Eastern chipmunk<br>*Tamias striatus* | ↓ | Pyare et al. 1993 |
| Short-tailed shrew<br>*Blarina brevicauda* | ↑ | Ford and Rodrigue 2001 |
| Smokey shrew<br>*Sorex fumeus* | 0↓ | Ford and Rodrigue 2001 |
| Masked shrew (Common shrew)<br>*Sorex cinereus* | 0↓ | Ford and Rodrigue 2001 |

The effects of disturbance on habitat structure caused by wildfire, clearcutting, forest harvest, and agriculture practices on small mammal occurrence and abundance are well studied and vary among species and habitat type (e.g. Ford and Rodrigue 2001, Klenner and Sullivan 2003, Burel et al. 2004, Fuller et al. 2004, Zwolak and Foresman 2008). In contrast, forest disturbance caused by invasive insects with consequence to small mammal community structure is mostly underrepresented in the literature. It seems that densities of habitat-specialists or rare species are negatively affected by different and varying degrees of disturbance (e.g. agriculture), but disturbance favors habitat-generalists densities (e.g. Burel et al. 2004). Other disturbances, such as fallow agriculture landscapes, can have a general positive role on small mammal species richness and biodiversity (e.g. Janova and Heroldova 2016). The degree in which habitat disturbance affects the abundance and community structure of small mammals is species-specific (review by Zwolak 2009). Herbivorous invasive insects, such as HWA, are known to create disturbed forests by alter nitrogen cycling and hydrologic processes (Brantley et al. 2015) and vegetation (Farnsworth et al. 2014) in the invaded forests, but how these changes impact associated animal communities remains unclear (review by Kenis et al. 2009)

Most of comparisons are based on uncontrolled natural experiments. Here, I take advantage of a replicated large-scale forest manipulation experiment to test for potential indirect effects of simulated hemlock woolly adelgid (HWA) invasion on forest microhabitats and small mammal community structure (Table 1). The purpose of this study was to 1) briefly describe microhabitat characteristics in disturbed forests that are generally known to influence small mammal distribution and 2) determine if simulated damage caused by hemlock woolly adelgid and preemptive logging impact small mammal communities.

## Methods

### Site Description

My work was conducted in north-central Petersham, Massachusetts, USA (42.47– 42.48°N, 72.22–72.21° W; elevation 215–300-m above sea level) within the Hemlock Removal Experiment at Harvard Forest. The Hemlock Removal Experiment is a replicated two-block (Ridge and Valley blocks) design with four ~90 × 90-m (~0.81-ha) canopy treatments plots (Hemlock, Logged, Girdled, and Hardwood). Two of the plots received canopy manipulations (Girdled and Logged) and the two plots that did not receive canopy manipulation and act as canopy controls (Hemlock and Hardwood). The canopy manipulations were applied in 2003 after baseline vegetation measurements were taken. For full detailed methodology on the experimental treatments, please refer to Ellison et al. (2010).

The experiment was conducted in hemlock dominated forests with similar topography (relative to their blocks) and aspects. The Girdled canopy manipulation was designed to simulate hemlock woolly adelgid (HWA) infestation. Physical damage to trees was applied by girdling all hemlock trees, excluding seedlings, with knives or chainsaws. Girdled hemlocks die within approximately 2 years after the treatment, but dead trees are left standing for several years’ post-mortem until they fall. This treatment mimics the response caused by HWA damage. Since the Girdled treatment was applied, the canopy density was reduced which resulted in a gradual increase of light availability to the understory vegetation over time (Farnsworth et al. 2012).

The Logged canopy manipulation was designed to mimic the effects of preemptive logging (as in many forest management plans) or commercial hemlock-salvage. All merchantable timber (hemlock, white pine, maple, birch, and oak) was harvested and removed. In contrast to the Girdled treatment, there was immediate light availability to the understory vegetation in the Logged plot (Farnsworth et al. 2012, Lustenhouwer et al. 2012) which allowed for early onset of vegetative growth of early sessional plants.

The Hemlock control plot was not manipulated and trees were intact to act as a control to Girdled and Logged treatments. However, since the experiment began, there have been sightings of the adelgid in the hemlock controls since 2009. Presently, there has been no reported hemlock mortality cause by the adelgid, but there has been increased light to the understory in Hemlock controls since 2009 (Kendrick et al. 2015) due to damage. The non-manipulated Hardwood plot represented the future of a Hemlock stand approximately 50 years after HWA invasion and preemptive logging.

### Sample Grid Layout

In 2012, I utilized a grid layout to examine the reduction of the eastern hemlocks on microhabitat structure and small mammal community dynamics. Sampling grids spanned 0.49-ha with sampling locations and trap stations placed 10-m apart in a 7 × 7 array within each of the two Hemlock, Hardwood, Girdled, and Logged plots (n=392). Grids were paced in a way to cover the most homogenous topography with the least amount of slope relief as possible.

### Microhabitat Characteristics

Microhabitat characteristics were derived from digital photographs of the ground and canopy of each trapping location taken during August 2013. One-m^2^ quadrates were placed over the trap location and then photographed to quantify small scale habitat characteristics that may affect small mammals. The camera (Canon EOS 7D, Canon Inc.) was placed approximately 1m from the ground to capture the entire 1m^2^ quadrat. Canopy photos were taken from the same position with the lens pointing to the canopy. Each ground photo (n=392) and each canopy photo (n=392) was labeled and scored using ImageJ (1.42q Java 1.6.0_version 10). Fifty points were randomly generated and overlaid on each digital photograph. The point location determined which characteristics would be scored. Ground and canopy characteristics that may be important to describe small mammal distribution included 1) rock, 2) soil, 3) woody debris, 4) leaf litter, 5) fungi, 6) vegetation, 7) open canopy, which was open sky or no canopy cover, 8) high canopy, which was characterized by canopy that was relatively far from the ground and considered old growth, and 9) low canopy, which was characterized by the canopy that was near the ground and considered new growth. Tree canopy was scored as “low” if the vegetation reached the photographers lens (approximately 1-m from the ground) and was scored “high” if the vegetation was greater than approximately 3-m and above. Each characteristic (i.e. rock, soil, vegetation, high canopy, etc…) for each sampling location (n=392) was calculated as a percent.

Randomized block analysis of variance (Randomized Block ANOVA) was used to determine significant difference of habitat characteristics between blocks (i.e. Ridge and Valley) and among treatments (i.e. Hemlock, Girdled, Logged, and Hardwood). Tukey’s Honest Significant Difference (HSD) post hoc pair-wise comparison test was used to identify differences between means that were greater than the expected standard error for the particular treatment (Tukey 1949, e.g. Gotelli and Ellison 2013).

### Small Mammal Live Trapping

Animals were captured in 2012 during summer months of June and July. Trapping was conducted during full, new, and half-moon conditions. Moonphase 3.3 (Tingstrom 2009) was used to determine the percentage of moon phase (illumination) for each trapping night. Traps were set for two consecutive nights in each block during similar peak moon phases. Sherman traps (H. B. Sherman, Tallahassee, FL USA)(9 × 9 × 3 inches) were placed within approximately 0.25-m of each sample location on the grid and traps set to sign. The goal was to promote captures, but not at the cost of assuming non-random captures by targeting specific species (see Hurlbert 1985). Sherman traps were used because they target the full spectrum of species in this assemblage. There is a bias towards capture of *Peromyscus spp*. when using Sherman traps (Dizney et al. 2008, Stephens and Anderson 2014), but the bias was the same among all treatments. Traps were baited with sunflower seeds to decrease trap disturbance by common regional predators (black bears, raccoons) and clean raw cotton was used for insulation. Traps were set about dusk and checked about dawn to limit sampling to nocturnal small mammals and to decrease stress caused by long term captivity. All traps were closed or folded down during the day.

Captured rodents and shrews were identified to species based on external morphology. Individual rodents were marked with colored non-toxic permanent ink. The color used was chosen based on the treatment where the induvial was captured. Therefore, individuals were not uniquely marked, but marks identified which treatment they were captured. Individuals were released at the same trapping location in which they were captured. All traps were closed or folded down during the day. All handling complied with rules and regulations set forth by the Animal Welfare Act and were approved by the Institutional Animal Care and Use Committee from University of Vermont (12-019) and Harvard University (12-04). Scientific collecting permit was obtained from the Massachusetts Department of Fish and Game (075.15SCM).

### Species Richness and Evenness

I used Chao1 (Chao 1984) abundance methods to estimate species richness among the treatments with 95% upper and lower confidence intervals. I used the shared abundance methods to estimate the number of shared species between the Hemlock control and the other treatments (Chao et al. 2000). The relative abundance or the proportion of the total assemblage that is represented by each species was calculated for each treatment. The average probability of interspecific encounter (PIE) was used to estimate species evenness for each treatment (Hurlbert 1971, Gotelli 2008). Confidence intervals for PIE were calculated from the standard deviations from the replicated treatments. Statistical support to accept the null hypotheses, which stated that there is no significant difference in species richness, shared species richness, and PIE among treatments, was determined by comparing the 95% confidence interval for the difference between the treatment means. If the difference in upper and lower confidence intervals of comparted groups did not contain zero, the null hypothesis was rejected (Knezevic 2008).

### Population Estimates with Mark-Recapture

Population of deer mice (*Peromyscus maniculatus*, Wanger 1845), white-footed mice (*Peromyscus leucopus*, Rafinewque 1818), southern red-backed voles (*Myodes gapperi*, Vigors 1830 [formerly *Clethrionomys gapperi*] were estimated with 10 nights of trapping in each block. Schnabel (1938) closed-population estimate was used due to the recapture marking methodology where animals were not marked as individuals. Confidence intervals (95% upper and lower) for estimated populations were calculated.

### Software and R Packages

All data were analyzed using R version 3.2.3 (R Core Team 2015). The package “reshape” version 0.8.5 (Wickham 2014) was used to restructure and aggregate data. The package “plyr” version 1.8.3 (Wickham 2015) was used for data frame manipulation. Packages lattice” version 0.20-33 (Sarkar 2015), “ggplot2” version 2.0.0 (Wickham and Chang 2015), and “grid” version 3.2.3 (Murrell 2005) were used for graphics. The package “agricolae” version 1.2.3 (de Meniburu 2015) was used for Tukey’s HSD grouping statistical procedures for microhabitat characteristics. The package “SpadeR” version 0.1.0 (Chao et al. 2015) was used to estimate species richness (‘ChaoShared’) and estimated shared species richness(‘ChaoShared’)

## Results

### Microhabitat Characteristics

There was no significant difference in the percent of rock cover among Hemlock (1.94%, SE= 0.39), Girdled (0.63%, SE= 0.33), Logged (1.04%, SE= 0.38), and Hardwood (1.29%, SE= 0.32) (F_3, 387_ = 2.412, P = 0.066), but there was a significant difference between Ridge and Valley blocks (F_1, 387_ = 22.34 P< 0.00001; Figure 1A). There was a significant difference in leaf litter ground cover among treatments (F_3, 387_ = 53.62, P<0.0001) and among blocks (F_1, 387_ = 20.23, P< 0.0001, Figure 1B). There was a difference in means among Hemlock (51.94%, SE= 1.90), Girdled (24.84%, SE= 1.78), and Logged (33.10%, SE= 1.81), but there was not a difference in means of percent leaf litter between Hemlock and Hardwood (51.29%, SE=2.06) (Figure 1B).

**Fig 1.**
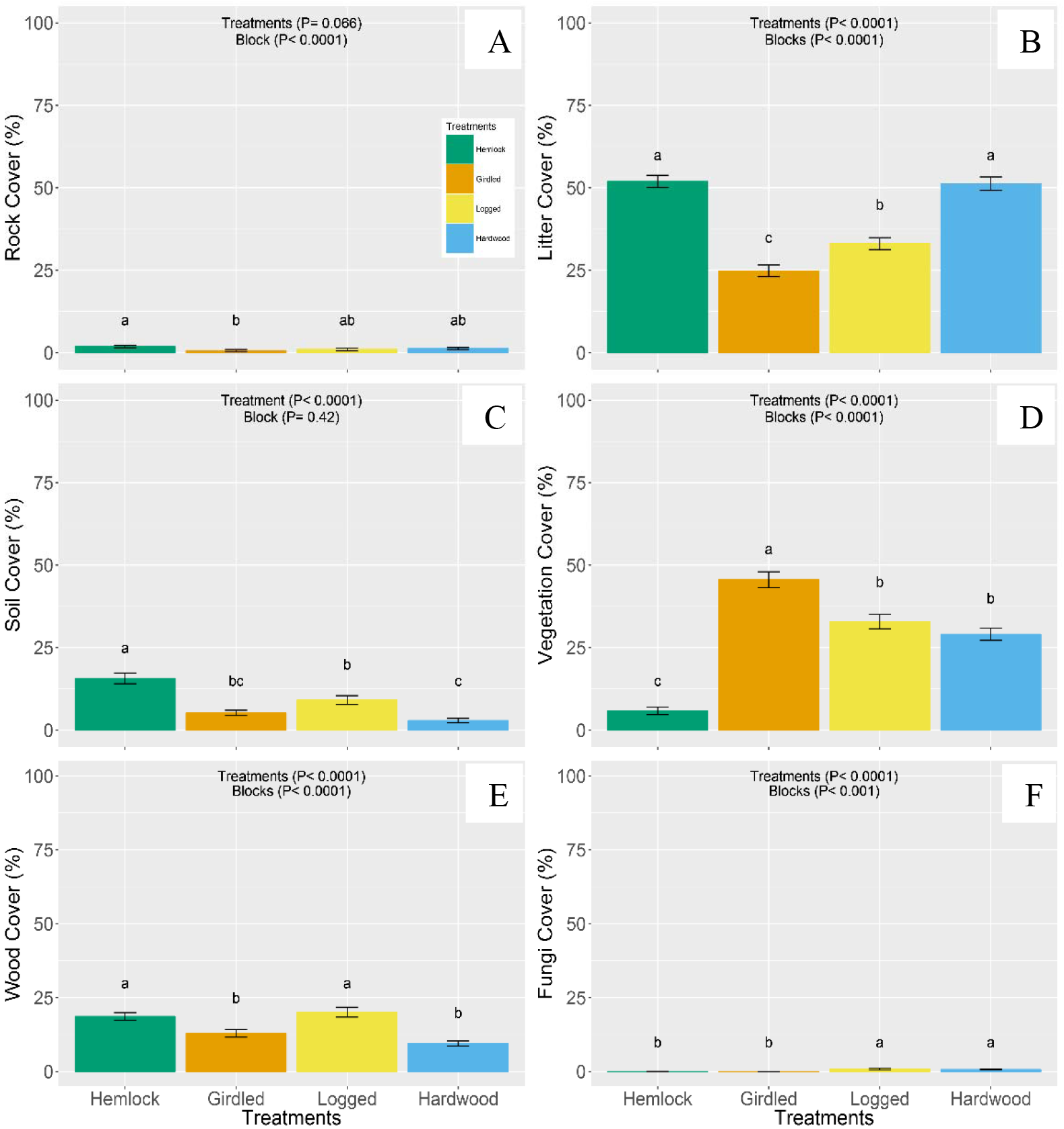
Mean (± SE) percent cover of microhabitat ground cover characteristics of rock (A), leaf litter (B), soil (C), vegetation (D), woody debris (E), and fungi (F) among Hemlock (green), Girdled (orange), Logged (yellow), and Hardwood (blue) treatments. Results of randomized block ANOVA for each characteristic indicated top-center of each graph. Lower case letters result of Tukey’s HSD grouping.

There was a significant difference in percent soil cover among treatments (F_3,387_ = 22.587, P< 0.0001), but not between blocks (F_1, 387_ = 0.651, P= 0.42, Figure 1C). There was a higher percent of soil cover on average in the Hemlock (15.69%, SE=1.63) treatments and the lowest percent of soil cover on average in the Hardwood (2.92%, SE= 0.66) treatment. There was no difference in soil means between the Girdled (5.22%, SE= 0.79) and Logged (9.12%, SE= 1.34) treatments (Figure 1C).

There was a significant difference in the percent of vegetation ground cover among treatments (F_3, 387_ = 76.06, P< 0.0001) and between blocks (F_1, 387_ = 22.93, P< 0.0001). There was a higher percent of understory vegetation in the Girdled (45.60%, SE= 2.39) treatment, but no difference between Logged (32.92%, SE= 2.18) and Hardwood (29.06%, SE= 1.85) treatments. Hemlock (5.86%, SE=1.15) treatments had the lowest percent vegetation cover among the treatments (Figure 1D). There was a significant difference in percent cover downed woody debris among treatments (F_3, 387_ = 15.68, P< 0.0001) and between blocks (F_1, 387_ = 29.42, P< 0.0001). There was a higher percent cover in Hemlock and Logged treatments, but there was no difference in the mean of woody debris cover between Hemlock (18.67%, SE= 1.26) and Logged (20.10%, SE=1.66). There was no difference between Girdled (12.92%, SE= 1.29) and Hardwood (9.51%, SE=0.83) means (Figure 1E). There was a significant difference in percent fungi cover among treatments (F_3, 387_ = 7.06, P< 0.0001) and between blocks, (F_1, 387_ = 8.036, P< 0.0001). Above fungi cover was low and there was no difference in the mean of fungi cover between Logged (0.88%, SE= 0.29) and Hardwood (0.67%, SE= 0.17) and no difference between Hemlock (0.02%, SE= 0.02) and Girdled (0.0%, SE= 0.0) (Figure 1F).

There was a significant difference in percent of open canopy among treatments (F_3, 387_ = 28.332, P< 0.0001), but not between blocks (F_1, 387_ = 3.33, P= 0.069). There was a difference in mean among Hemlock (8.22%, SE= 0.49), Girdled (23.14%, SE= 1.52), Logged (19.65%, SE= 1.82), and Hardwood (13.51%, SE= 0.62), but not between Girdled and Logged treatments (Figure 2A). There was a significant difference in percent of high canopy cover among treatments (F_3, 387_ = 169.02, P< 0.0001), but not between blocks (F_1, 387_ = 1.56, P= 0.21). There was not a significantly higher percent in high canopy between Hemlock (86.41%, SE= 1.08) and Hardwood (79.55%, SE= 1.73) treatments, but there was a difference high canopy cover in Girdled (37.49%, SE= 2.97) and Logged (24.47%, SE=3.04) treatments with Logged having the least amount of high canopy cover (Figure 2B). There was a significant difference in low canopy cover among treatments (F_3, 387_ = 85.12, P< 0.0001), but not between block (F_1, 387_ = 3.77, P=0.05). The Logged (55.84%, SE= 3.59) treatment had a higher percent canopy cover than Girdled (39.31%, SE= 3.57), Hemlock (5.37%, SE= 0.92), and Hardwood (6.90%, SE= 1.66). However, there was not a difference in low canopy percent cover between Hemlock and Hardwood (Figure 2C).

**Fig 2.**
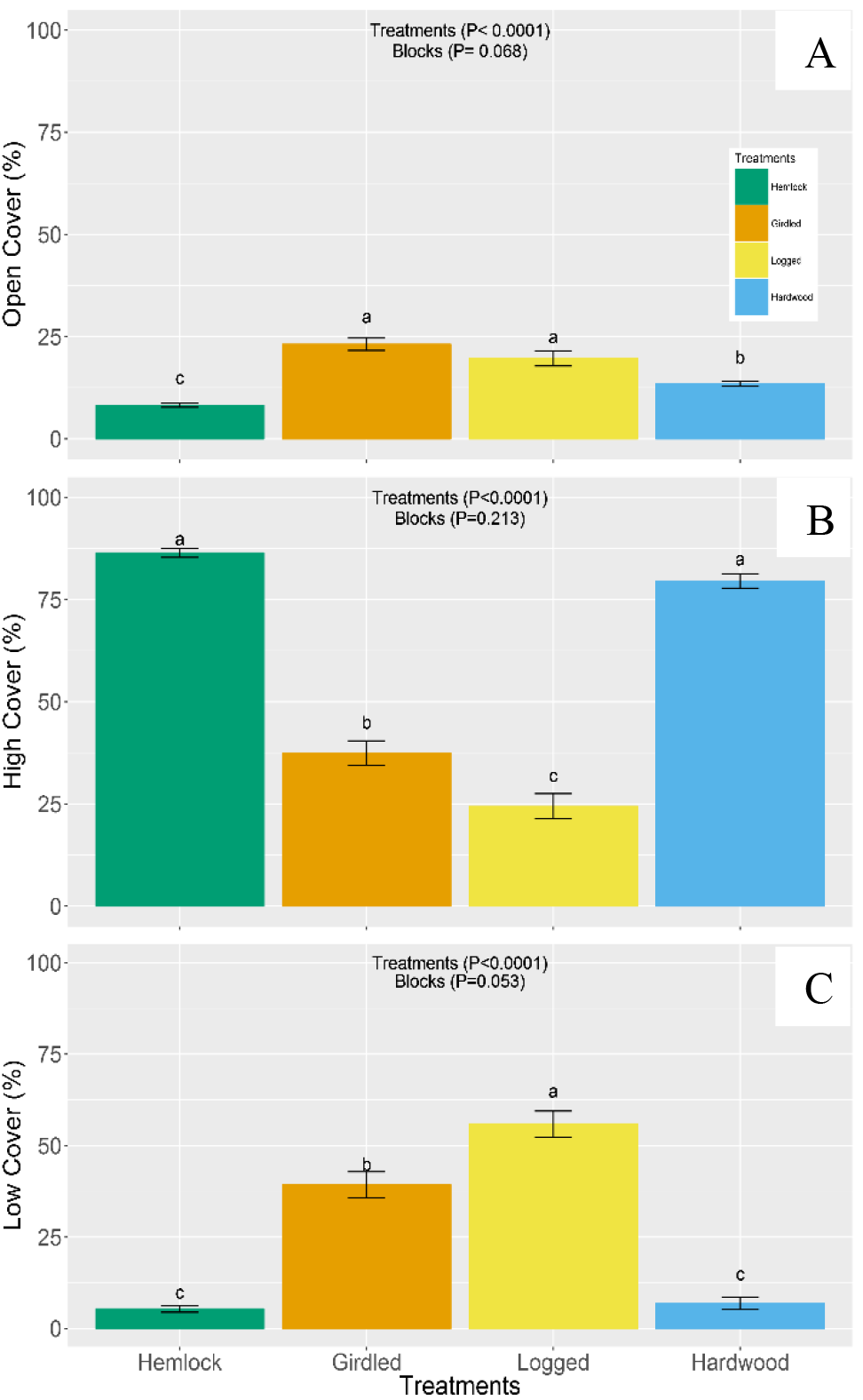
Mean (± SE) percent cover of microhabitat canopy cover characteristics open canopy (A), high canopy (B), and low canopy (C) among Hemlock (green), Girdled (orange), Logged (yellow), and Hardwood (blue) treatments. Results of randomized block ANOVA for each characteristic indicated top-center of each graph. Lower case letters above treatment are result of Tukey’s HSD grouping.

### Small Mammal Captures

There were 4,131 trapping nights and 18.7% capture success among all treatments. I trapped in the Ridge block for 12 nights and in the Valley for 10 nights. There were 2,183 traps set in the Ridge block and trapping success varied among Hemlock (17%), Girdled (22%), Logged (16%), and Hardwood (22%) treatments. There were fewer traps set in the Ridge-Hardwood treatment (n=420) than other treatments in the Ridge block (n=588) due to a change in property management midway through the season. In the Valley block, there were 1,948 traps set and trapping success varied among Hemlock (14%), Girdled (20%), Logged (15%), and Hardwood (24%) treatments. Although there was a slight difference in the number of traps used in the Ridge-Hardwood plot than in the Valley-Hardwood plot, the percent trapping success was comparable.

### Species Richness and Evenness

The observed small mammal species (i.e. rodents and shrews) varied slightly among treatments (Figure 3). There were more species found in the Girdled (8) than in the Logged (7), Hemlock (6), and Hardwood (6) treatments. Deer mice, white-footed, southern red-backed, and short-tailed shrews (*Blarina brevicauda*, Gray 1838) were found among all treatments (Figure 3). Southern flying squirrels (*Glaucomys Volans*, Linnaeus 1758) were most abundant in the control plots, but one was captured in a Girdled plot (Figure 3) and only on one occasion. Eastern chipmunks (*Tamias striatus*, Linnaeus 1758) and masked shrews (*Sorex cinereus*, Kerr 1792) were more abundant in disturbed treatments than in controls (Figure 3). Eastern chipmunks were not captured in the Hemlock controls and masked shews were not captured in the Hardwood controls (Figure 3). Woodland jumping mice (*Napaeozapus insignis*, Miller 1891) and woodland voles (*Microtus pinetorum*, LeConte 1830) were only captured in the disturbed treatments and with very low captures (Figure 3). Relative capture abundance for deer mice ranked highest in Hemlock, Logged, and Hardwood and southern red-backed vole abundance ranked highest in the Girdled treatment (Figure 3). While there was a difference in the assemble of small mammals (not all species captured among all treatments), there was no significant difference in community evenness. The average PIE (F_3, 387_= 0.34, P=0.79) among Hemlock (PIE= 0.59, lower 95%CI= 0.14, upper 95%CI= 1.00), Girdled (PIE= 0.63, lower 95%CI= 0.63, upper 95%CI= 0.74), Logged (PIE= 0.68, lower 95%CI=0.63, upper 95%CI= 0.73), and Hardwood (PIE= 0.63, lower 95%CI= 0.58, upper 95%CI= 0.68) were not significantly different among treatments (F_1, 387_= 2.29, P= 0.22).

**Fig 3.**
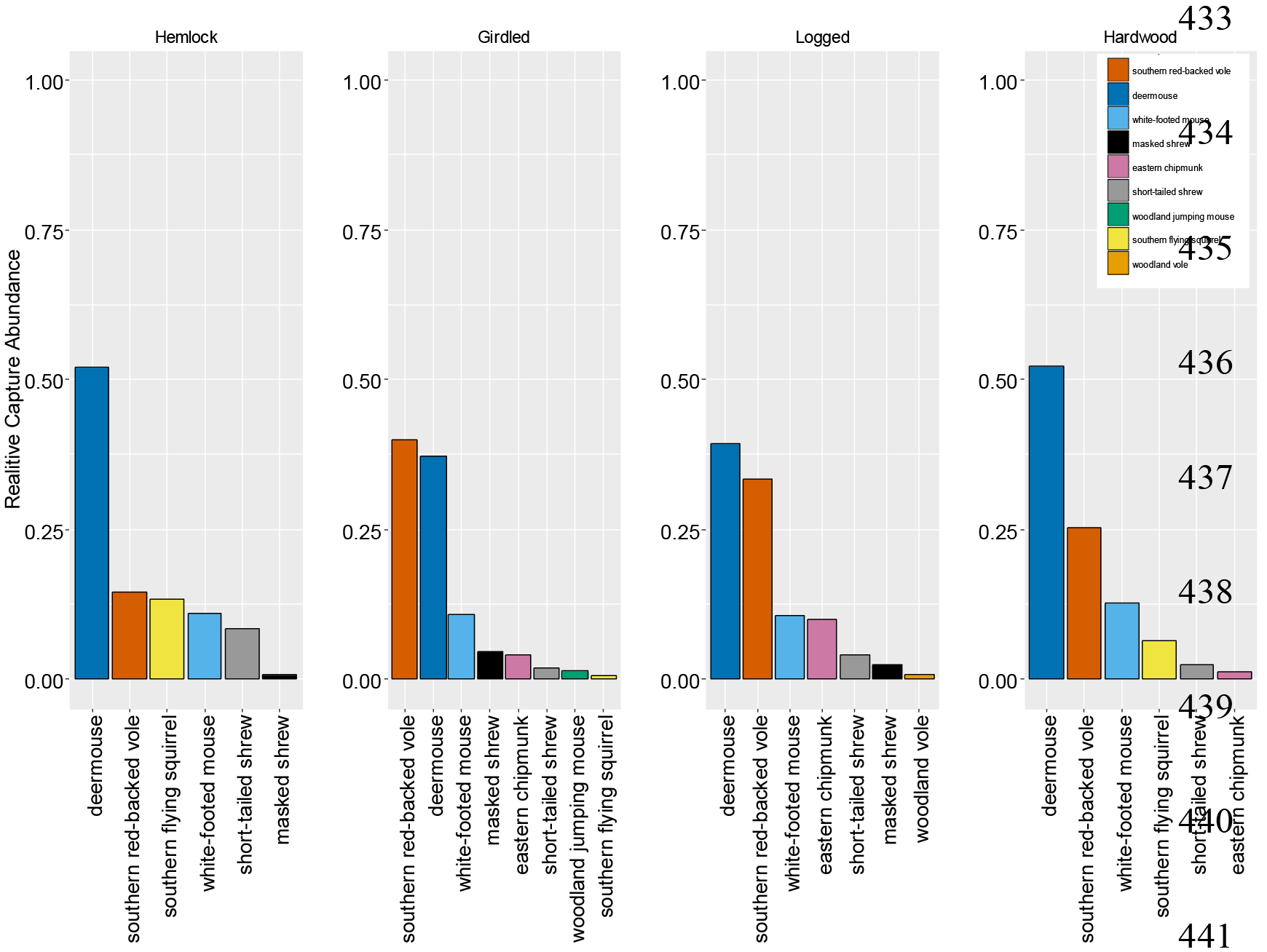
Rank relative abundance graph of small mammals in 2012 among canopy treatments (left to right): Hemlock, Girdled, Logged, and Hardwood). Each bar and each color represents a different species. The height of the bar is the relative abundance of the species in each treatment.

The estimated species richness was highest in the Girdled treatment (n= 8, lower 95%CI = 8.07, upper 95%CI = 9.59, Figure 4), followed by the Logged treatment (n= 7, lower 95%CI = 7.0, upper 95%CI = 8.45 Figure 4). The estimated species richness was the same (n=6) in the Hemlock (lower 95%CI = 6.0, upper 95%CI = 7.40) and in Hardwood controls (lower 95%CI = 6.0, upper 95%CI = 6.49, Figure 4). There was a significant difference in estimate species richness between the controls and Girdled treatment, but not between controls and Logged treatment (Figure 4). There was no significant difference between Hemlock and Hardwood controls and between Girdled and Logged treatments (Figure 4). There were six estimated shared species between Hemlock and Girdled (SE= 0.57, lower 95%CI = 5.35, upper 95%CI = 7.85), five shared between Hemlock and Logged (SE= 0.46, lower 95%CI = 4.43, upper 95%CI = 6.29) and six shared between Hemlock and Hardwood (SE= 0.0, lower 95%CI = 5.0, upper 95%CI = 5.0)

**Fig 4.**
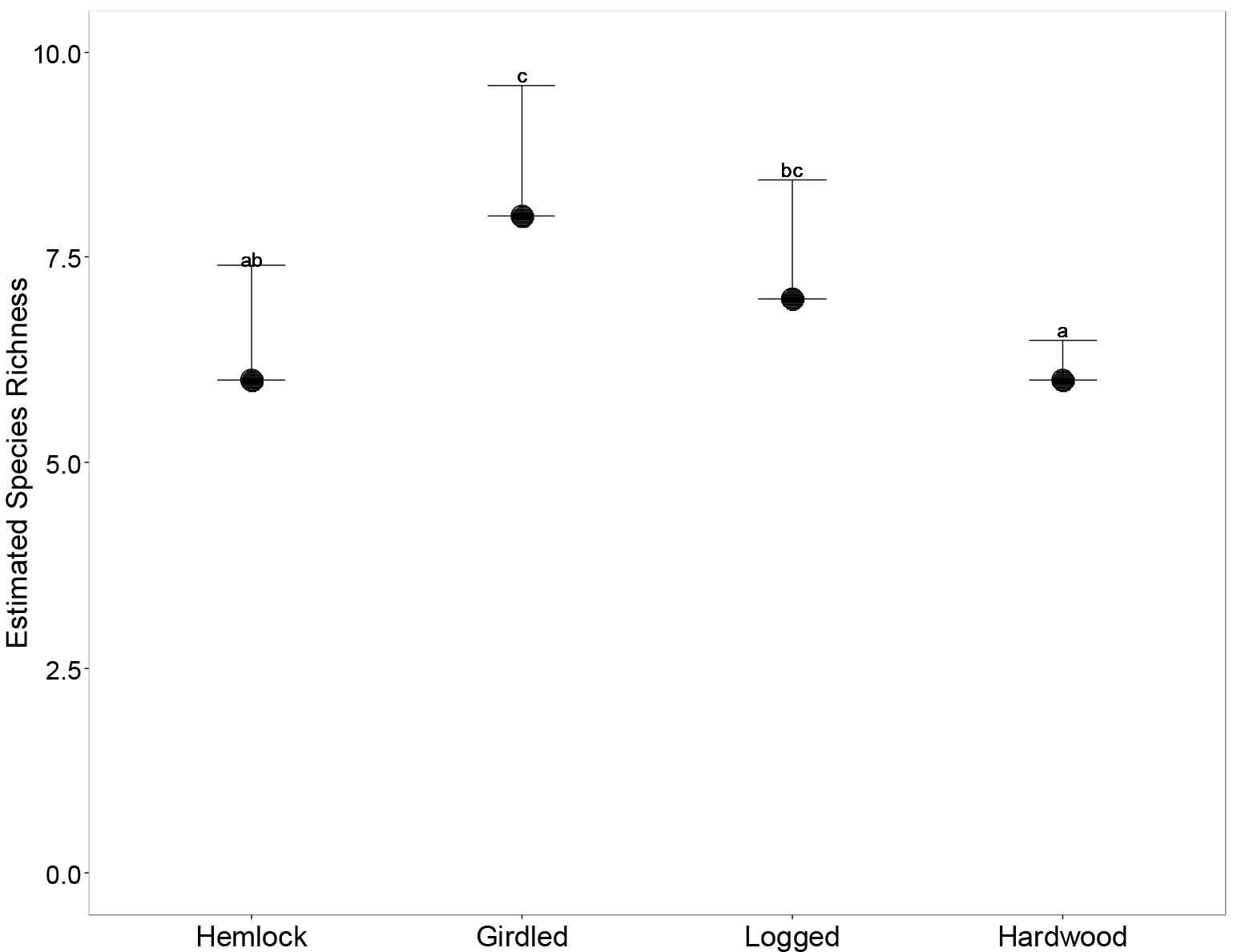
Chao1 estimated species richness (dots) with lower and upper 95% confidence intervals (error bars) for Hemlock, Girdled, Logged, and Hardwood canopy treatments. Letters indicate groupings based on CI overlap where different letters indicate significantly different groups. The estimated species richness in Hemlock (a) control differs significantly from Girdled (c), but not from Logged (b) and Hardwood (a). Girdled treatments (c) differs significantly from Hardwood (a), but not Logged (c).

### Population Estimates

There was a denser population of deer mice in the Logged treatment (N-hat = 40.7 per 0.64ha, lower 95%CI = 27.17, upper 95%CI = 64.33) than in the Hemlock control (N-hat = 17.14 per 0.64ha, lower 95%CI = 13.19, upper 95%CI = 24.47, Figure 5), but all other treatments do not have overlapping error bars (Knezevic 2008). The southern red-backed vole population was denser in the Girdled (N-hat = 84.4 per 0.64ha, lower 95%CI = 59.80, upper 95%CI = 136.41) and Logged treatments (N-hat = 47.11 per 0.64ha, lower 95%CI = 31.15, upper 95%CI = 85.62) than the Hemlock (N-hat = 8.14per 0.64ha, lower 95%CI =4.85, upper 95%CI = 14.89) and Hardwood controls (N-hat= 17.2 per 0.64ha, lower 95%CI = 12.73, upper 95%CI = 25.43, Figure 5). There was no difference in population density of white-footed mice among Hemlock (N-hat= 9.0 per 0.64ha, lower 95%CI = 4.74, upper 95%CI = 19.68), Girdled (N-hat= 8.31 per 0.64ha, lower 95%CI = 4.85, upper 95%CI = 15.60), Logged (N-hat= 10.87 per 0.64ha, lower 95%CI = 6.03, upper 95%CI = 30.91) and Hardwood treatments (N-hat= 7.20 per 0.64ha, lower 95%CI = 4.84, upper 95%CI = 11.25, Figure 5).

## Discussion

I found small scale microhabitat characteristics (Figures 1 and 2), small mammal community assemblage (Figure 3), estimated species richness (Figure 4), and southern red-backed vole populations (Figure 5B) were affected by girdled and logged disturbance, but community evenness (PIE) and mice populations (Figure 5A & C) were not.

**Fig 5.**
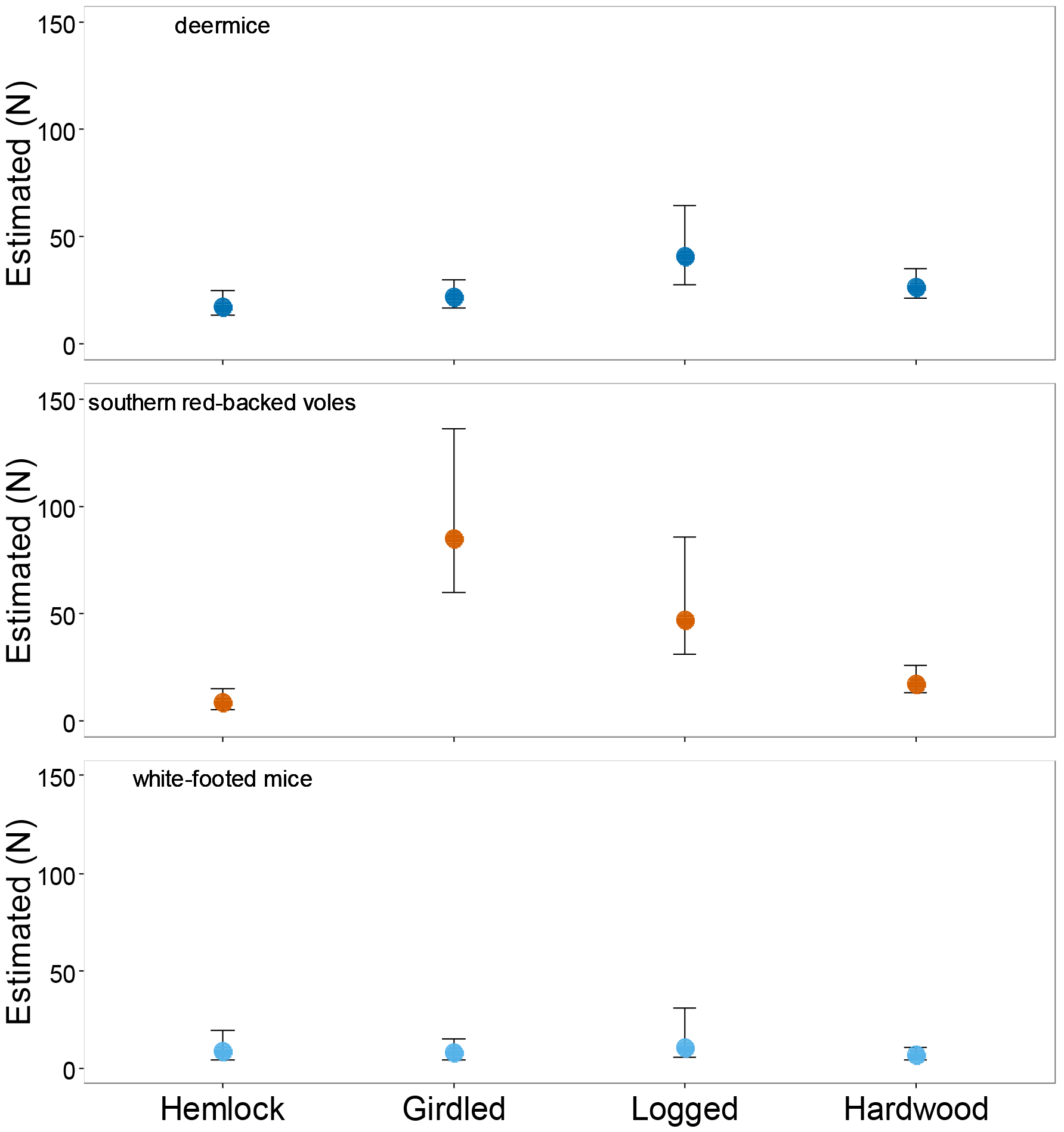
Schnebel estimated population (N-hat) with lower and upper 95% confidence intervals (error bars) for Hemlock, Girdled, Logged, and Hardwood canopy treatments for deer mice, southern red-backed voles, and white-footed mice (top to bottom).

Hemlock woolly adelgid and logging increased the percent ground cover of vegetation, but decreased the percent ground cover of leaf litter relative to hemlock controls. These disturbances also deceased the amount of high canopy cover which allowed for an increase in open canopy cover and low canopy cover. Heterogeneous changes in habitat structure caused by invasive species can create patches of suitable and unsuitable habitat. These variations in habitat or patch quality may influence the site occupancy (the probability that a particular species is present at a site; MacKenzie et al. 2002). While these disturbances may seem minor, they can have detrimental effects on small mammal distribution, especially habitat specialists.

Overall, estimated species richness did increase in the Girdled treatment relative to the Hemlock control (Figure 4) and there were more species represented in the Girdled treatment than in the Hemlock control (Figure 3). However, not all species that were sampled were found in the Girdled treatment and several were rarely captured (Figure 3). For example, southern flying squirrels were not captured in logged treatments at all and only one was captured in the girdled treatment. This suggests that the presence or site occupancy of southern flying squirrels may decrease as hemlock woolly adelgid continues to spread and destroy hemlock forests in New England and southern flying squirrels will depend more on hardwood forests in the future. Given that no southern flying squirrels were found in the logged treatments it seems safe to assume that preemptive logging management would be equally devastating to these arboreal rodents as girdling from hemlock woolly adelgid damage. Although northern flying squirrels were not captured in this study, I predict that their populations would also decrease dramatically as adelgid spreads northward. Unlike southern flying squirrels that utilize both hemlock and hardwood stands (primarily hardwood), northern flying squirrels (*Glaucomys sabrinus*, Shaw 1801) depend on old growth forests (Ransome and Sullivan 1997) such as old eastern hemlock forests. If the spread of the adelgid continues to increase northward as it is predicted, the northern flying squirrels may not have time to adapt to the changing forests and the species could be lost.

Community evenness of small mammals did not differ among treatments. This could be due to the large variation of PIE estimates in the Hemlock controls. When PIE was separated by block, there was a significant difference between PIE estimates in Valley (PIE = 0.43) and Ridge (PIE = 0.76) Hemlock blocks. This suggests that slight changes in the landscape (elevation and slope) may influence species distribution or at least influence capture ability of some animals. Regardless of the community evenness, there were differences in the overall community assemblage and species richness estimates between Girdled treatment and controls. Although deer mice and white-footed mice populations were not affected by girdled and logged treatments, the southern red-backed vole populations were positively affected by the disturbances (Figure 5). It seems that habitat generalists (e.g. deer mice, white-footed mice) may not be as impacted by hemlock woolly adelgid and logging as habitat specialists (e.g. southern flying squirrels), which would support previous studies (e.g. Millán de la Peña et al. 2003, Burel et al. 2004, Zowlak 2009, Janova and Heroldova 2016). Finding larger population densities of southern red-backed voles within disturbed areas of old-hemlock forests support findings that these voles are not necessarily old-forest specialists, but are more associated with habitat features such as woody debris (Keinath and Hayward 2003) or well decayed woody debris (Fauteix et al. 2012).

## Conclusion

The loss of this foundation species did impact microhabitat characteristics, small mammal communities, and alter species richness enough to warrant caution in discussing the severity of eastern hemlock population decline. The local increase in small mammal species richness, represented by a snap-shot short-term study, should not be interpreted as HWA being “good” for small mammal diversity. On the contrary, these data suggest that there are varying degrees in which small mammals will be impacted with continued spread of hemlock woolly adelgid and destructive management practices (preemptive logging). In addition to the loss of eastern hemlocks, the implications for regional changes in small mammal communities may have dramatic consequential effects on forest dynamics as each small mammal species provide ecosystem functions across a large environmental gradient including, 1) the increase of forest range with the dispersal of seed (e.g. Steele et al. 2006; Beck and Vander Wall 2010; Yu et al. 2013), 2) being used as bio-indicators for forest health (e.g. Haim and Izhaki 1994; Pearce and Veiner 2005; Leis et al. 2008), 3) determination of seed fate through seed foraging, dispersal, caching, and hoarding behaviors (e.g. Steele et al. 2011), and 4) facilitating vegetation growth in forests and fields (Ostfeld et al. 1993; Howe et al. 2006). They are also important food resources for many vertebrates (e.g. Sullivan et al. 2004, Sundell et al. 2013) and are hosts to diverse groups of parasites (e.g. Vandergrift et al. 2009, Kuhnen et al. 2011). We need to consider the ramification of eastern hemlock loss on other species and how those species contribute to the overall ecosystem function.

## Acknowledgements

I thank undergraduate researchers Elizabeth Kennett, Emma Cornin, and Jefferson Franca de Jesus for their hard work in and out of the field. I thank Chris Degrassi for his programing assistance and technical support. The manuscript benefited from input from Nick Gotelli, Alison Brody, Aaron Ellison, Bill Kilpatrick, and Becca Rowe.

## References

Beck MJ, Wall SBV (2010) Seed dispersal by scatter-hoarding rodents in arid environments. J Eco 98:1300–1309

Burel F, Butet A, Delettre Y, Millàn de la Peña N (2004) Differential response of selected taxa to landscape context and agricultural intensification. Landscape and Urban Planning 67:195–204

Chao A (1984) Nonparametric estimation of the number of classes in a population. Scandinavian J Statistics 11:65–270

Chao A, Hwang W-H, Chen Y-C, Kuo C-Y (2000) Estimating the number of shared species in two communities. Statistica Sinica 10:227–246

de Meniburu Fd (2016) Package ‘agricolae’ version 1.2-4. Statistical Procedures for Ag Res. CRAN:1–157

Dizney L, Jones PD, Ruedas LA (2008) Efficacy of three types of live traps used for surveying small mammals in the Pacific Northwest. Northwestern Naturalist 89:171–180

Ellison AM, et al. ( 2005a) Loss of foundation species: Consequences for the structure and dynamics of forested ecosystems. Frontiers in Eco and the Environ 3:479–486

Ellison AM, Chen J, Burnham CK, Díaz D, Lau M. (2005b) Changes in ant community structure and composition associated with hemlock decline in New England. Third Symposium on Hemlock Woolly Adelgid: 280–289

Ellison AM, Barker Plotkin AA, Foster DR, Orwig DA (2010) Experimentally testing the role of foundation species in forests: the Harvard Forest Hemlock Removal Experiment. Methods in Eco & Evo 1:168–179

Farnsworth EJ, Barker Plotkin AA, Ellison AM (2012) The relative contributions of seed bank, seed rain, and understory vegetation dynamics to the reorganization of *Tsuga canadensis* forests after loss due to logging or simulated attack by *Adelges tsugae*. Can J For Res 42:2090–2105

Ford WM, Rodrigue JL (2001) Sorcid abundance in partial overstory removal harvests and riparian area in industrial forest landscapes of central Appalachians. For Eco Manage 152:159–168

Foster DR, Orwig DA (2006) Preemptive and salvage harvesting of New England forests: When doing nothing is a viable alternative. Con Bio 20:959–970

Fuller AK, Harrison DJ, Lachowski HJ (2004) Stand scale effects of partial harvesting and clearcutting on small mammals and forest structure. For Eco and Manage 191:373–386

Gotelli NJ (2008) Gotelli, N.J. 2008. A primer of ecology, fourth edition. Sinauer, Sunderland, Massachusetts

Gotelli NJ, Ellison AM (2013) A primer of ecological statistics: Second edition. Sinauer Associates, Inc.

Graves S, Maldonado J, Wolff JO (1988) Use of ground and arboreal microhabiatats by *Peromyscus leucopus* and *Peromyscus maniculatus*. Can J Zool 66:277–278

Henein K, Wegner J, Merriam G (1998) Population effects of landscape model manipulation on two behaviourally different woodland small mammals. Oikos 81:168–186

Howe HF, Zorn-Arnold B, Suillivan A, Brown JS (2006) Massive and distinctive effects of meadow voles on grassland vegetation. Ecology 87:3007–3013

Hurlbert SH (1984) Pseudoreplication and the design of ecological field experiments. Ecological Monographs 54:187–211

Janovaa E, Heroldova M (2016) Response of small mammals to variable agricultural landscapes in Central Europe. Mammalian Bio 81:488–493

Keinath, DA, and Hayward, GD (2003) Red-backed vole (*Clethrionomys gapperi*) response to distrubance in subalpine forests: Use of regenerating patches. J Mamm 84:956–966

Kendrick JA, Ribbons RR, Classen AT, Ellison AM (2015) Changes in canopy structure and ant assemblages affect soil ecosystem variables as a foundation species declines. Ecosphere 6:1–21

Kenis M, Augerr-Rozenberg MA, Roques AR, Timms L, Pe’re’ C, Cock MJW, Settele J, Augustin S, Lopez-Vaamonde C (2009) Ecological effects of invasive alien insects. Biol Invasions (2009) 11:21–45

Kizlinski ML, Orwig DA, Cobb RC, Foster DR (2002) Direct and indirect ecosystem consequences of an invasive pest on forests dominated by eastern hemlock. J Biogeogr 29:1489–1503

Klenner W, Sullivan TP (2003) Partial and clear-cut harvesting of high-elevation spruce–fir forests: implications for small mammal communities. Can J For Res 33:2283–2296

Knezevic A (2008) StatNews #73. Overlapping confidence intervals and statistical significance. Cornell University: Cornell Statistical Consulting Unit. Report no.

Kuhnen VV, Graipel ME, Pinto CJC (2011) Differences in richness and composition of gastrointestinal parasites of small rodents (Cricetidae, Rodentia) in a continental and insular area of the Atlantic Forest in Santa Catarina state, Brazil. Brazil J Bio 72:563–567

Lustenhouwer MN, Nicoll L, Ellison AM (2012) Microclimate effect of the loss of a foundation species from New England forest. Ecosphere 3:1–16

MacKenzie DI, Nichols JD, Lachman GB, Droege S, Royle JA, Langtimm CA (2002) Estimating site occupancy rates when detection probabilities are less than one. Ecology 83:2248–2255

McClure MS (1989) Importance of weather to the distribution and abundance of introduced adelgid and scale insects. Ag and For Meterology 47:291–302

McClure MS (1991) Density-dependent feedback and popualation cycles in *Adelges tsugae* (Homoptera: Adelgidae) on *Tsuga canadensis*. Environ Entomol 20:258–264

Murrell P (2005) Package’grid’ R Graphics Chapman & Hall/CRC Press

Orwig DA, Cobb RC, D’Amato AW, Kizlinski ML, Foster DR (2008) Multi-year ecosystem response to hemlock woolly adelgid infestation in southern New England forests. Can J For Res 38:834–843

Orwig DA, Foster DR (1998) Forest response to the introduced hemlock woolly adelgid in southern New England, USA. J Torrey Bot Soc 125:60–73

Orwig DA, Foster DR, Mausel DL (2002) Landscape patterns of hemlock decline in New England due to the introduced hemlock woolly adelgid. J Biogeogr 29:1475–1487

Ostfeld RS, Canham CD (1993) Effects of meadow vole population density on tree seedling survival in old fields. Ecology 74:1792–1801

Pyare S, Kent JA, Noxon DL, Murphy MT (1993) Acorn preference and habitat use in easter chipmunks. Am Midland Nat 130:179–183

Ransome DB, Sullivan TP (1997) Food limitation and habitat preference of *Glaucomys sabrinus* and *Tamiasciurus hudsonicus*. J Mamm 78:538–549

Sarkar D (2015) Package ‘lattice’. CRAN:1–157

Schnabel ZE (1938) The estimation of total fish population of a lake. The American Mathematical Monthly 45:348–352

Siddig AAH, Ellison AM, Mathewson BG (2016) Assessing the impacts of the decline of *Tsuga canadensis* stands on two amphibian species in a New England forest. Ecosphere 7(11):e01574. 10.1002/ecs2.1574

Steele MA, Bugdal M, Yuana A, Bartlowa A, Buzalewski J, Lichti N, Swihart R (2011) Cache placement, pilfering, and a recovery advantage in a seed-dispersing rodent: Could predation of scatter hoarders contribute to seedling establishment? Acta Oecologica 37:554–560

Steele MA, Manierre S, Genna T, Contreras TA, Smallwood PD, Pereira ME (2006) The innate basis of food-hoarding decisions in grey squirrels: evidence for behavioural adaptations to the oaks. Animal Behaviour 71:155–160

Stephens, RB and Anderson EM (2014) Effects of trap type on small mammal richness, diversity, and mortality. Wildl Soc Bull 38:619–627

Sullivan TP, Sullivan DS, Reid DG, Leung MC (2004) Weasels, voles, and tress: Influence of mustelid semiochemicals on vole populations and feeding damage. Ecol App 14:999–1015

Sundell J, Hara RBO, Helle P, Hellstedt P, Henttonen H, Pietiäinen H (2013) Numerical response of small mustelids to vole abundance: delayed or not? Oikos 122:1112–1120

Taulam JF, Seaman DE (2000) Assessung southern flying squirrel, *Glaucomys volans*, habitat selection with kernel home range estimation and GIS. Can Field-Naturalist 114:591–600

Tingley MW, Orwig DA, Field R, Motzkin G (2002) Avian response to removal of a forest dominant: consequences of hemlock woolly adelgid infestations. J of Biogeogr, 29: 1505–1516

Tingstrom H (2009) Moonphase 3.3- the Northern Hemisphere version

Tukey JW (1949) Comparing individual means in the analysis of variance. Biometrics 5:99–114

Vickery WL, Rivest D (1992). The influence of weather on habitat use by small mammals. Ecography 15:205–211

Wickham H (2015) Package ‘reshape’. Pages 1–20. CRAN

Wickham H (2016) Package ‘plyr’. Pages 1–62. CRAN

Wickham H, Chang W (2016) Package ‘ggplot2’. Pages 1–197. CRAN

Wolff JO, Schauber EM, Edge WD (1997) Effects of habitat loss and fragmentation on the behavior and demography of gray-tailed voles. Con Bio 11:945–956

Yu F, Wang D, Shi X, Yi X, Li G (2013) Seed dispersal by small rodents favors oak over pine regeneration in the pine-oak forests of the Qinling mountains, China. Scandinavian J Res 28:540–549

Zwolak R (2009) A meta-analysis of the effects of wildfire, clearcutting, and partial harvest on the abundance of North American small mammals. For Eco Manage 258:539–545

Zwolak R, Foresman KR (2008) Deer mouse demography in burned and unburned forest: no evidence for source-sink dynamics. Can J Zool 86:83–91

